# First and broad detection of three typical carbapenemase genes on the surfaces of commercially available spices worldwide and isolation of complete NDM-1 genes from black pepper, cumin, and clove

**DOI:** 10.1101/2020.06.09.141614

**Authors:** Minako Mochizuki, Sumio Maeda

**Author notes:** Correspondence Sumio Maeda, Department of Food Science and Nutrition, Nara Women’s University Graduate School of Humanities and Sciences, Kitauoya-nishimachi, Nara, Nara 630-8506, Japan.

## Abstract

The spread of multidrug-resistant bacteria, particularly those producing carbapenemases, has become a major public health concern. The presence of carbapenemase genes has primarily been reported in clinical samples, whereas the presence of these genes in commercially available foods has insufficiently been studied despite its growing importance. The present study aimed to detect and characterize carbapenemase genes (*bla*_NDM,_ *bla*_IMP_, *bla*_KPC_, and *bla*_OXA-48-like_) on the surfaces of commercially available spices using PCR to amplify conserved regions of these genes. It was revealed that DNAs of these genes are commonly present (13 genes/29 samples) on spices derived from at least 9 different countries. This is the first detection of any carbapenemase gene on eight spices (black pepper, cumin, clove, cardamom, mustard, caraway, parsley, and rosemary) among these. This is also the first detection of the *bla*_IMP_ and *bla*_NDM_ as well as the broad detection of the *bla*_OXA-48-like_ on spices. We also isolated complete, functional *bla*_NDM-1_ genes from three spices (black pepper, cumin, and clove) up to the present.

## INTRODUCTION

Antimicrobial resistance (AMR) has become great concern worldwide in the clinical field as well as in the field of food production and distribution, or the food chain (1–7). Carbapenem resistance is one of the most severe forms of AMR (1,8), and carbapenemase [a subgroup of β-lactamase (bla)]-producing bacteria, such as carbapenemase-producing Enterobacteriaceae, are spreading worldwide (9,10). Although the spread of carbapenemase genes has mainly been reported in clinical samples to date (1), several recent studies have reported the presence of carbapenemase-producing bacteria in primary industry products and environments (3–7,11,12); it is considered that this increasing presence at such settings is owing to the routine use of antibiotics for livestock, poultry, and crops (13,14) as well as occasionally owing to contamination via workers or organic fertilizers (15–17). Among those studies, a few have described the presence of carbapenemase genes in certain commercially available foods (5–7). However, there is insufficient knowledge regarding the status of contamination and spread of those genes on commercially available foods, particularly those distributed worldwide via international trade.

In the current study, we attempted to detect carbapenemase genes from commercially available spices. Spices are often produced in countries where carbapenemase-producing bacteria are spreading, and are exported worldwide. Because almost all spice products are distributed as dried forms to the market, fewer precautions have been taken against bacterial presence in spices compared with those taken against bacterial presence in perishable foods. Even in dried spices, dormant bacterial cells including endospores and viable but non-culturable cells (18–20) can survive for long periods. Despite these situations, there is only one study on the presence of a carbapenemase gene on the surface of a spice; Zurfluh, et al. (7) reported that a carbapenemase gene *bla*_OXA-181_ was present on commercially available coriander. In the present study, to investigate the presence of four typical carbapenemase genes [*bla*_IMP_ (IMP), *bla*_NDM_ (NDM), *bla*_KPC_ (KPC), and *bla*_OXA-48-like_ (OXA-48)] on various spices, we performed PCR analysis on 29 samples of 13 species. We describe the first detection of the IMP and NDM genes as well as the broad detection of the OXA-48 gene on the surfaces of commercially available spices obtained from various countries worldwide. Moreover, we isolated three full-length NDM-1 genes from three spice samples.

## RESULTS

Spice samples in sealed bottles that are listed in Table 1 were purchased at grocery stores in Japan between October 2017 and November 2018. PCR assay using the primers for the conserved regions of each carbapenemase gene was conducted for the DNA purified from the surface of spice samples (29 samples of 13 species). One PCR reaction contained template DNA derived from mg order of each spice sample (the column of “mg of spice” in Table 1); such amounts of spices are within the range of daily usage of most spices. The details of PCR experiments are described in MATERIALS AND METHODS and the Supporting Information (Table S1, and Fig. S1). A total of 13 PCR products were identified as parts of independent carbapenemase genes (Table 1). Among 29 samples tested, 12 samples were carbapenemase sequence-positive. Among 13 spice species, carbapenemase sequence-positive products were detected in 9 species (black pepper, cumin, mustard, coriander, cardamom, caraway, parsley, rosemary, and clove). These spices were derived from at least 9 different countries worldwide (Malaysia, India, Canada, Morocco, Guatemala, Egypt, France, the United States, Indonesia, Sri Lanka, and one unspecified).

**Table 1.** Results of PCR, sequencing, and BLAST search of carbapenemase gene fragments derived from spice samples

| sample |  |  |  | iden-<br>tified<br>gene | acce-<br>ssion<br>No. | results of BLAST search |  |  |  |  |
| --- | --- | --- | --- | --- | --- | --- | --- | --- | --- | --- |
| spice<br>source | No. | *mg<br>of<br>spice | country |  |  | blastn |  |  | blastp |  |
|  |  |  |  |  |  | hit<br>subject | bp/bp | iden-<br>tity<br>(%) | aa/aa | iden-<br>tity<br>(%) |
| black<br>pepper | 1 | 4.0 | Malaysia | IMP | LC48<br>4335 | IMP-like | 183/191 | 96 | 63/63 | 100 |
|  | 2 | 4.0 | unknown | NDM | LC48<br>4332 | NDM-1, -2,<br>-4, -6, -7,<br>-8, -9, -11,<br>-12, -14,<br>-15, -16,<br>-18, -19,<br>-22, -25,<br>-28 | 115/115 | 100 | 45/45 | 100 |
| cumin | 1 | 4.0 | India | NDM | LC48<br>4333 | NDM-1, -2,<br>-4, -6, -7,<br>-8, -9, -11,<br>-12, -14,<br>-15, -16,<br>-18, -19,<br>-22, -25,<br>-28 | 115/115 | 100 | 45/45 | 100 |
|  |  |  |  | OXA-48 | LC48<br>4337 | OXA-48,<br>-48b, -252,<br>-514,<br>-48-like | 192/192 | 100 | 63/63 | 100 |
|  | 2 | 4.0 | Turkey | - | - | - | - | - | - | - |
| sweet<br>pepper | 1 | 4.7 | China | - | - | - | - | - | - | - |
|  | 2 | 3.1 | South<br>Korea | - | - | - | - | - | - | - |
| mustard | 1 | 2.5 | Canada | OXA-48 | LC48<br>4344 | OXA-48,<br>-48b,<br>-48-like | 191/192 | 99 | 63/63 | 100 |
|  | 2 | 16 | Canada | - | - | - | - | - | - | - |
|  | 3 | 7.0 | Canada | - | - | - | - | - | - | - |
| coriander<br>(seed) | 1 | 2.3 | Morocco | OXA-48 | LC48<br>4340 | OXA-48,<br>-48b, -252,<br>-514,<br>-48-like | 192/192 | 100 | 63/63 | 100 |
|  | 2 | 1.0 | Morocco | - | - | - | - | - | - | - |
| cardamom | 1 | 9.4 | Guate-<br>mala | IMP | LC48<br>4336 | IMP-like | 181/191 | 95 | 61/63 | 96 |
|  | 2 | 2.3 | India | - | - | - | - | - | - | - |
| caraway | 1 | 6.6 | Nether-<br>lands | - | - | - | - | - | - | - |
|  | 2 | 5.7 | Nether-<br>lands | - | - | - | - | - | - | - |
|  | 3 | 3.5 | Egypt | OXA-48 | LC48<br>4341 | OXA-48,<br>-48b, -252,<br>-514,<br>-48-like | 192/192 | 100 | 63/63 | 100 |
| parsley | 1 | 1.3 | France | OXA-48 | LC48<br>4338 | OXA-48b,<br>-199,<br>-48-like | 192/192 | 100 | 63/63 | 100 |
|  | 2 | 0.6 | France | - | - | - | - | - | - | - |
|  | 3 | 3.4 | United<br>States | OXA-48 | LC48<br>4339 | OXA-48b,<br>-199,<br>-48-like | 192/192 | 100 | 63/63 | 100 |
| rosemary | 1 | 3.7 | Morocco | OXA-48 | LC48<br>4343 | OXA-48,<br>-48b, -252,<br>-514,<br>-48-like | 191/192 | 99 | 63/63 | 100 |
|  | 2 | 3.1 | Albania | - | - | - | - | - | - | - |
| clove | 1 | 1.1 | Indonesia | OXA-48 | LC48<br>4342 | OXA-48,<br>-48b, -252,<br>-514,<br>-48-like | 192/192 | 100 | 63/63 | 100 |
|  | 2 | 2.2 | Sri Lanka | NDM | LC48<br>4334 | NDM-1, -2,<br>-4, -6, -7,<br>-8, -9, -11,<br>-12, -14,<br>-15, -16,<br>-18, -19,<br>-22, -25,<br>-28 | 115/115 | 100 | 45/45 | 100 |
| basil | 1 | 0.6 | Egypt | - | - | - | - | - | - | - |
|  | 2 | 1.4 | Egypt | - | - | - | - | - | - | - |
| sichuan<br>pepper | 1 | 4.0 | China | - | - | - | - | - | - | - |
|  | 2 | 3.0 | China | - | - | - | - | - | - | - |
| oregano | 1 | 0.7 | India | - | - | - | - | - | - | - |
|  | 2 | 0.9 | Turkey | - | - | - | - | - | - | - |

-: not detected or not applicable; aa: amino acid
\* mg of spice: mg of spice sample, from which template DNA per PCR reaction was derived.

Furthermore, IMP sequences were detected in two samples (black pepper and cardamom), NDM sequences were detected in three samples (black pepper, cumin, and clove), and OXA-48 sequences were detected in eight samples [cumin, mustard, coriander, caraway, parsley (2 samples), rosemary, and clove]. No KPC sequences were detected in any sample. Although most samples contained only one kind of carbapenemase sequence, cumin sample No. 1 contained two kinds of carbapenemase sequences (NDM and OXA-48).

Of the identified 13 sequences, 9 were 100% identical to the known sequences; however, four sequences (IMP genes of black pepper No. 1 and cardamom No. 1; OXA-48 genes of mustard No. 1 and rosemary No. 1) exhibited 95%–99% identities in the nucleotide sequence (“bp/bp” in Table 1) and 96%–100% in the amino acid sequence (“aa/aa” in Table 1) without containing any frame-shift or nonsense mutation, suggesting that they are candidates for parts of novel gene variants.

Among the 13 carbapenemase sequence-positive samples described above, we successfully isolated and identified three full-length NDM genes from black pepper No. 2, cumin No.1, and clove No. 2 (Table 1) at this point. They were cloned into expression vectors; subsequently, these plasmids were introduced to *E. coli* cells. All these cells resulted in the expression of ampicillin and meropenem resistance, while the vector alone did not express such resistance (the data of black pepper are shown in Fig. 1 as a representative). Therefore, these genes were demonstrated to function as carbapenemase genes. According to BLAST analysis, it was revealed that all these genes (accession number LC507187, LC532142, and LC532143) were 100% identical to NDM-1 (accession number NG049326.1) (21,22).

**Fig. 1.**
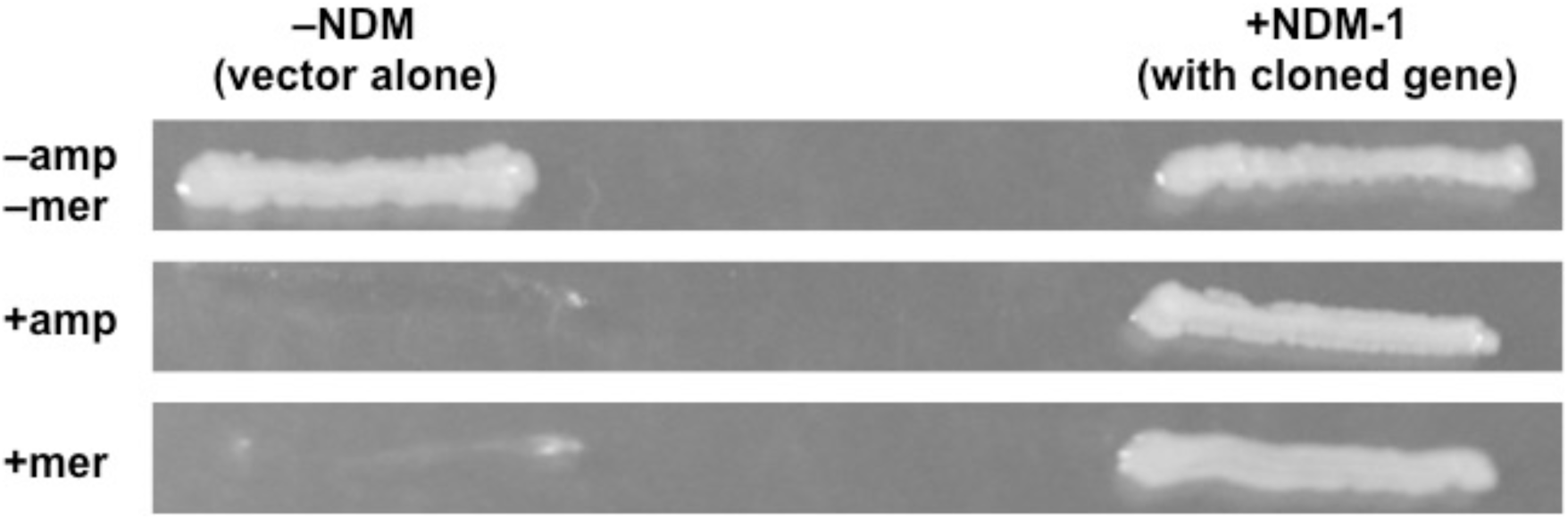
Expression of carbapenemase activity from the NDM-1 gene isolated from black pepper No. 2. Carbapenemase activity of the NDM-1 gene isolated from black pepper No. 2 was examined by growing *E. coli* JM83 cells expressing this gene on an expression vector pHSG299 [sample “+NDM-1 (with cloned gene)”] on LB agar in the absence (–amp, –mer) or presence of beta-lactam antibiotics [+amp: ampicillin (50 µg/mL); +mer: meropenem (0.25 µg/mL)]. Sample “–NDM (vector alone)” is the negative control of JM83 harboring the original pHSG299 without any insert.

## DISCUSSION

In the present study, we identified 13 partial sequences of carbapenemase genes from 29 spice samples (13 species). To the best of our knowledge, this is the first detection of IMP and NDM genes on the surfaces of commercially available spices. In all spices (black pepper, cumin, clove, cardamom, mustard, caraway, parsley, and rosemary) except coriander, this is the first detection of carbapenemase genes. Moreover, we detected the OXA-48 gene in a broad range of samples [cumin, mustard, coriander, caraway, parsley (2 samples), rosemary, and clove]. In addition, we succeessfully isolated complete sequences of the NDM-1 gene from three spices (black pepper, cumin, and clove).

Because these spices were derived from at least 9 different countries worldwide, our results suggest that the contamination of spices with carbapenemase genes, or possibly with carbapenemase-producing bacteria, is spreading worldwide; this finding is consistent with those of the studies conducted on other foods (5–7). In the present study, NDM genes were detected in the samples derived from India, Sri Lanka, and unknown country; the former two countries are consistent with the origin of this gene (9,21,22). Moreover, OXA-48 gene was detected in the samples derived from eight different countries, which is consistent with the fact that OXA-48 genes are broadly distributed worldwide (9,23,24). Four identified sequences of two IMP genes and two OXA-48 genes are suggested to be parts of new gene variants, implying that the food chain may be a blind route for the occurrence and spread of novel carbapenemase variants.

In Japan, IMP is the major carbapenemase that has been detected in clinical samples (10,25), whereas the presence of other carbapenemases remains minor despite their gradual increase in recent years (26–31). Therefore, detection of NDM genes, especially of the complete NDM-1 gene, in spices that are popularly available in Japan would strongly impact the public health in Japan. Because this situation is probably not limited to Japan but is valid for other countries as well, further detailed and broader surveys on the presence of carbapenemases in the global food chain are required, as recently suggested by the World Health Organization (1,14).

We successfully isolated full-length, functional NDM-1 genes directly from three independent spice samples using the PCR-based method. It should be noted that even if such functional genes exist as “silent” naked DNA on the surfaces of spices, they have potentialities to be absorbed by the encountering bacteria via transformation (32,33), subsequently rendering the DNA functional and replicable in living cells. Further studies, including isolation of the other full-length genes from spice samples and elucidation of their existence within or outside bacteria, are required to fully comprehend our results.

In conclusion, we report the first detection of the IMP and NDM genes, including the complete NDM-1 gene, as well as the broad detection of the OXA-48 gene on the surfaces of commercially available spices. Moreover, in most spices except coriander, this is the first detection of any carbapenemase gene. Because of a rather high rate detection of the carbapenemase genes on the spices worldwide, our present study demonstrates the spread of AMR via the global food chain. Further studies would provide newer insights into this critical public health issue.

## MATERIALS AND METHODS

### Materials

Spice samples in sealed bottles that are listed in Table 1 were purchased at grocery stores in Japan between October 2017 and November 2018. Phosphate-buffered saline (PBS; 1.47 mM potassium phosphate monobasic; 8.1 mM sodium phosphate bibasic; 2.7 mM potassium chloride; and 137 mM sodium chloride; pH 7.4 at 25°C), ampicillin, and kanamycin were obtained from Sigma (St. Louis, MO, US). The synthetic PCR primers used (Table S1) were obtained from Eurofins Genomics (Tokyo, Japan). KAPA Taq Extra HotStart ReadyMix was obtained from KAPA Biosystems (Woburn, MA, US). EmeraldAmp® PCR MAX Master Mix and agarose were obtained from Takara Bio (Kusatsu, Japan). Whole genome amplification (WGA) kit (REPLI-g® Mini Kit) was obtained from Qiagen (Venlo, Netherlands). Tryptic soy broth (TSB) was obtained from Becton, Dickinson (Franklin Lakes, NJ, US). Chelex100 Resin was obtained from Bio-Rad (Hercules, CA, US). Distilled water (DNase- and RNase-free, molecular biology grade) was obtained from Merck Millipore (Burlington, MA, US). Meropenem, agar powder (guaranteed reagent grade), and other general reagents were obtained from Wako (Osaka, Japan). An *E. coli* strain JM83 (*rpsL, ara, Δ(lac-proAB), Φ80dlacZΔM15*) (34) and a kanamycin-resistant cloning vector pHSG299 (35) were obtained from the National BioResource Project: *E. coli* (NIG, Japan) (https://shigen.nig.ac.jp/ecoli/strain/).

### Preparation of template DNA

Template DNA for PCR assay was purified from the surface of spices as follows. Each spice sample (1.7–22 g) was washed with 50 mL of fresh PBS. Following the filtration of the resultant PBS solution to remove residue, total DNA was purified from the solution using proteinase K and phenol/chloroform (36), resulting in 0.8–3.5 mL of template DNA/H_2_O solution for each sample. Moreover, Chelex treatment (37) was performed for some samples before the PCR assay.

### PCR analysis of universal bacterial DNA (16S-rRNA)

To confirm the presence of bacterial DNA in the template DNA samples, the presence of bacterial 16S-rRNA gene was examined using PCR. PCR assays using the universal bacterial 16S-rRNA primers (each 0.25 µM; Table S1) and template DNA (0.5–3.0 µL/10µL PCR solution) were performed as follows: an initial denaturation at 94°C for 5 min, 35 cycles of 30 s at 94°C, 30 s at 56°C, 30 s at 72°C, and final extension at 72°C for 7 min. Consequently, samples that showed positive results were used for the subsequent PCR analysis for carbapenemase genes.

### Preparation of positive control DNA for carbapenemase genes

To determine whether the carbapenemase gene primers can function under the reaction conditions containing sample DNA solutions prepared from spices, we prepared artificial positive control DNAs that could be amplified with the carbapenemase gene primers but yielded different sized PCR products. These control DNAs were prepared with sequence-adding PCR using pUC19 as the original template, in which universal primers (M4 and RV), an original primer (Table S1), and primers with extended sequences corresponding to carbapenemase primers were used (Table S1).

### PCR, sequencing, and data analysis for carbapenemase genes

PCR for four carbapenemase genes (IMP, NDM, KPC, and OXA-48) was conducted using primers detailed in Table S1 and template DNA [0.5–3.0 µL (normally 1.0 µL) in 10µL of the reaction mixture] in the presence or absence of the corresponding positive control DNA. PCR assays using KAPA Taq Extra HotStart ReadyMix were performed as follows: an initial denaturation at 94°C for 5 min, 50 cycles of 30 s at 94°C, 30 s at 52°C, 30 s at 72°C, and a final extension at 72°C for 7 min. Further, PCR assays using EmeraldAmp® PCR MAX Master Mix for detection of IMP and NDM gene were performed with an initial denaturation at 94°C for 5 min, 16 cycles of 30 s at 94°C, 30 s at 55.0°C –50.0°C (−0.3°C per cycle), 30 s at 72°C, 34 cycles of 30 s at 94°C, 30 s at 50°C, 30 s at 72°C, and a final extension at 72°C for 7 min. For detection of OXA-48 and KPC, PCR assays were performed with an initial denaturation at 94°C for 5 min, 10 cycles of 30 s at 94°C, 30 s at 60.5°C –55.5°C (−0.5°C per cycle), 30 s at 72°C, 40 cycles of 30 s at 94°C, 30 s at 55.5°C, 30 s at 72°C, and a final extension at 72°C for 7 min. The amplified products were analyzed using conventional 2.8% agarose-gel electrophoresis. Some of the DNA fragments in the gels were cut out, purified, and used for an additional PCR assay to amplify those products. The re-amplified PCR products obtained were sequenced by Macrogen Japan (Tokyo, Japan), and the data were analyzed using BLAST (http://blast.ncbi.nlm.nih.gov/Blast.cgi). The gene names of the highest hit score in each sample are shown in Table 1.

### Isolation, cloning, and expression of full-length NDM gene obtained from black pepper, cumin, and clove

Full-length NDM genes were isolated from the DNA of black pepper No. 2, cumin No.1, and clove No. 2 using WGA and subsequent PCR using primers including those for N-terminal and C-terminal ends of the genes (Table S1). These genes were cloned into the Pst I-BamH I site of pHSG299, transformed to and expressed in *E. coli* JM83. The expression of carbapenemase activity was examined in LB broth or on LB agar containing 50 µg/mL ampicillin or 0.25 µg/mL meropenem. The sequences of the cloned genes were read and analyzed as described above.

## DISCROUSURE

There are no conflicts of interest to declare.

## ACKNOWLEDGEMENTS

We appreciate Mariya Ueda, Shiori Nakaoka, Miyuki Nakatani (Nara Women’s University) for their technical assistance. We are also grateful to Enago (www.enago.jp) for English editing and proofreading services.

## List of Abbreviations

AMR: Antimicrobial resistance;
bla: β-lactamase;
IMP: *bla*_IMP_;
KPC: *bla*_KPC_;
NDM: *bla*_NDM_;
OXA-48: *bla*_OXA-48-like_.

## SUPPLEMENTAL MATERIAL

Table S1. Primer list

Fig. S1. Flow of preparation of positive control DNAs for IMP, OXA, KPC, and NDM genes.

## DATA AVAILABILITY

The DNA sequences with accession numbers that we reposited in DDBJ/ENA/Genbank will be released after acceptance of manuscript.

