## Supplemental Table S1 & Figure S1 for "First and broad detection of three typical carbapenemase genes on the surfaces of commercially available spices worldwide and isolation of complete NDM-1 genes from black pepper, cumin, and clove"

Table S1. Pimer list

| PCR product | Primer name | Primer sequence(5'-3') | Source | Explanation |
|---|---|---|---|---|
| Bacterial 16S-rRNA fragment | Bacteria Universal (B)-F | GAG TTT GAT CMT GGC TCA G | ref. S1 | for detection of 16S-rRNA |
| | Bacteria Universal (B)-R | GTA TTA CCG CGG CKG CTG | | |
| Positive control of IMP | IMP-FR | GGA ATA GAG TGG CTT AAY TCT CCA GGA AAC AGC TAT GAC | this study | for preperation and amplification of positive control DNA |
| | IMP-RF | GGT TTA AYA AAA CAA CCA CCG TTT TCC CAG TCA CGA C | this study | |
| Positive control of OXA | OXA-FR | TTC GGC CAC GGA GCA AAT CAG CAG GAA ACA GCT ATG AC | this study | |
| | OXA-RF | GAT GTG GGC ATA TCC ATA TTC ATC GCA GTT TTC CCA GTC ACG AC | this study | |
| Positive control of KPC | KPC-FF | CCT TCA TGC GCT CTA TCG TTT TCC CAG TCA CGA C | this study | |
| | KPC-RR | TTT GTA AGC TTT CCG TCA CGC AGG AAA CAG CTA TGA C | this study | |
| Positive control of NDM | R-lacZ-F | CGC CAT TCG CCA TTC AG | this study | |
| | NDM-FR | GGC AGC ACA CTT CCT ATC TCA GGA AAC AGC TAT GAC | this study | |
| | NDM-RF | GTT GAT CTC CTG CTT GAT CCG CCA TTC GCC ATT CAG | this study | |
| IMP fragment | IMP-F | GGA ATA GAG TGG CTT AAY TCT C | ref. S2 | for detection of carbapenemase gene fragments |
| | IMP-R | GGT TTA AYA AAA CAA CCA CC | | |
| NDM fragment | NDM-F | GGC AGC ACA CTT CCT ATC TC | ref. S3 | |
| | NDM-R | GTT GAT CTC CTG CTT GAT CC | | |
| KPC fragment | KPC-F | CCT TCA TGC GCT CTA TCG | ref. S4 | |
| | KPC-R | TTT GTA AGC TTT CCG TCA CG | | |
| OXA fragment | OXA-F | TTC GGC CAC GGA GCA AAT CAG | ref. S5 | |
| | OXA-R | GAT GTG GGC ATA TCC ATA TTC ATC GCA | | |
| NDM full-sequence | NDM-F(complete) | ATG GAA TTG CCC AAT ATT ATG C | this study | for detection of full-length NDM gene |
| | NDM-R(compelete) | TCA GCG CAG CTT GTC G | this study | |
| NDM full-sequence for cloning | Pst I +NDM-F(complete) | GCC TGC AGG ATG GAA TTG CCC AAT ATT ATG C | this study | for cloning of full-length NDM gene |
| | BamH I +NDM-R(compelete) | GCG GAT CCT CTC AGC GCA GCT TGT CG | this study | |

First and broad detection of three typical carbapenemase genes on the surfaces of commercially available spices worldwide and isolation of complete NDM-1 genes from black pepper, cumin, and clove

Mochizuki and Maeda

Supplemental Material

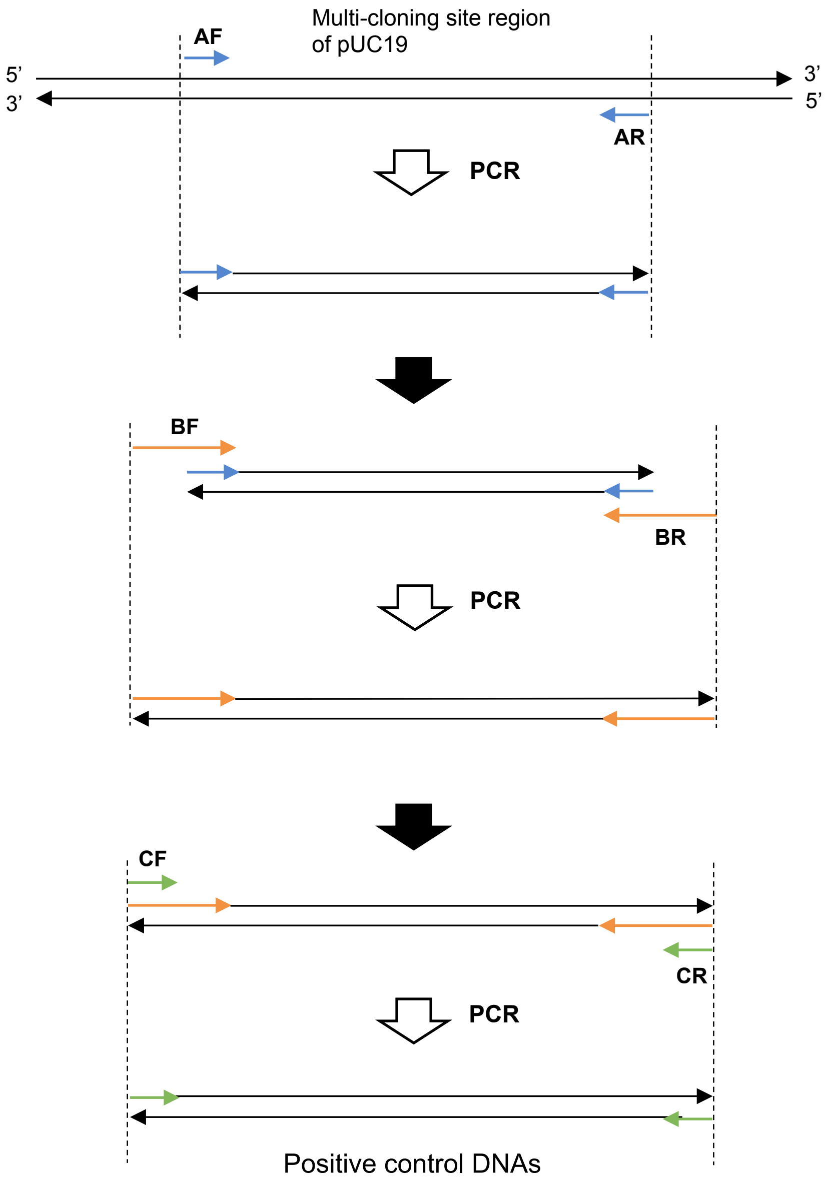


**Fig. S1 Flow of preparation of positive control DNAs for IMP, OXA, KPC, and NDM genes.**
To prepare each positive control DNA, the following primers (listed in Table S1) were used.
For IMP, AF: M13M4, AR: M13RV, BF: IMP-RF, BR: IMP-FR, CF: IMP-R, and CR: IMP-F.
For OXA, AF: M13M4, AR: M13RV, BF: OXA-RF, BR: OXA-FR, CF: OXA-R, and CR: OXA-F.
For KPC, AF: M13M4, AR: M13RV, BF: KPC-FF, BR: KPC-RR, CF: KPC-F, and CR: KPC-R.
For NDM, AF: R-lacZ-F, AR: M13RV, BF: NDM-RF, BR: NDM-FR, CF: NDM-R, and CR: NDM-F.
